# LncRNA SChLAP1 promotes cancer cell proliferation and invasion via its distinct structural domains and conserved regions

**DOI:** 10.1101/2025.01.28.635288

**Authors:** Mihyun Oh, Roshni Nagesh Kadam, Zahra Sadruddin Charania, Srinivas Somarowthu

## Abstract

Long non-coding RNAs (lncRNAs) play key roles in a range of biological processes and disease progression. Despite their functional significance and therapeutic potential, lncRNAs’ mechanisms of action remain understudied. One such lncRNA is the Second Chromosome Locus Associated with Prostate-1 (SChLAP1). SChLAP1 is overexpressed in malignant prostate cancer and is associated with unfavorable patient outcomes, such as metastasis and increased mortality. In this study, we demonstrated that SChLAP1 possesses distinct structural domains and conserved regions that may contribute to its function. We determined the secondary structure of SChLAP1 using chemical probing methods combined with mutational profiling (DMS-MaP and SHAPE-MaP). Our *in vitro* secondary structural model revealed that SChLAP1 consists of two distinct secondary-structural modules located at its 5’ and 3’ ends, both featuring regions with a high degree of structural organization. Our *in vivo* chemical probing identified potential protein-binding hotspots within the two modules. Overexpression of the modules led to a notable increase in cancer cell proliferation and invasion, proving their functional significance on the oncogenicity of SChLAP1. In conclusion, we discovered functionally important, independent modules with well-defined structures of SChLAP1. These results will serve as a guide to explore the detailed molecular mechanisms by which SChLAP1 promotes aggressive prostate cancer, ultimately contributing to the development of SChLAP1 as a novel therapeutic target.

## INTRODUCTION

While the majority of the human genome is transcribed into RNA through a highly regulated process, protein-coding genes represent less than 2% of the total transcripts (1,2). This has prompted the exploration of the remaining non-coding transcripts, and recent advances in sequencing technologies have uncovered a large number of long non-coding RNAs (lncRNAs) (1). LncRNAs are RNA molecules longer than 200 nucleotides that lack protein-coding capabilities. Similar to protein-coding messenger RNAs (mRNAs), most lncRNAs are transcribed by RNA polymerase II and undergo extensive processing, including splicing, capping, and polyadenylation (2,3). Additionally, lncRNAs display distinct patterns of expression specific to cell types, developmental stages, and subcellular localization, suggesting intrinsic functional significance (4,5).

LncRNAs are increasingly recognized for their functional importance and potential therapeutic applications. Recent research indicates that lncRNAs are involved in crucial processes such as cell development, aging, and disease progression (6). Furthermore, several lncRNAs have shown promise as biomarkers for cancer progression; for example, the lncRNA PCA3 has been FDA-approved for the early detection and management of prostate cancer (7,8). While lncRNAs are garnering significant interest as potential therapeutic targets against various diseases, including cancer, the mechanism by which lncRNAs exert their functions remains largely unexplored. To date, only a limited number of lncRNAs, such as Xist, have been extensively studied, despite an estimated 20,000 lncRNAs being annotated in the human genome (9).

Investigating the structure-function relationships of lncRNAs can provide insights into their underlying molecular mechanisms (10). LncRNAs present exceptional structural complexity, and this structural complexity of lncRNAs allows them to mediate various biological functions via interactions with proteins and nucleic acids (11). Emerging studies suggest that lncRNAs’ functions were more dependent on structural conservation than on primary sequence conservation (12,13). For instance, the tumor suppressor MEG3 has conserved, structured domains. Mutations that disrupt the long-range, pseudoknot interaction between its two distal structural motifs destroy MEG3-dependent p53 stimulation, whereas mutations that retain the same secondary structure remain functional (14). An emerging theme from lncRNA structural studies is that lncRNAs have modular architectures, with their modular domains mediating distinct functional roles, exemplified by lncRNAs GAS5 and MUNC (15–18).

This study focuses on the lncRNA Second Chromosome Locus Associated with Prostate-1 (SChLAP1), which is overexpressed in malignant prostate cancer, and it relates to poor patient outcomes, including metastasis and a high mortality rate (19). Since it is already considered to be a useful biomarker for predicting metastatic progression in prostate cancer, SChLAP1 is of interest as a promising therapeutic target (7,19). However, the molecular mechanism behind SChLAP1’s oncogenic role in cancer development is not well understood. Thus, further research into the oncogenic mechanism of SChLAP1 is urgently needed.

As a step towards a comprehensive understanding of SChLAP1’s molecular mechanism, we characterized its secondary structure using chemical probing approaches. First, we determined the *in vitro* secondary structure model of SChLAP1, demonstrating that SChLAP1 forms a complex secondary structure. Specifically, we identified structurally stable domains, 5’ and 3’ ends of SChLAP1, which also exhibit evolutionary conservation. Finally, we used overexpression analysis to show that these domains independently promote prostate cancer cell invasion and proliferation.

## MATERIAL AND METHODS

### Cell culture

LNCaP and 22Rv1 cells (kind gifts from the lab of Dr. Alessandro Fatatis, Drexel University College of Medicine, Philadelphia, PA, USA) were cultured in RPMI 1640 media (Gibco, Cat # 11875085) supplemented with 10% FBS (Cytiva, Cat # SH30541.03) and 1% penicillin/streptomycin antibiotic (Genesee Scientific, Cat # 25-512). Cells were incubated at 37°C and 5% CO_2_ and maintained by splitting at 80-90% confluence. For LNCaP cells, 40μM cell strainers (Thermo Fisher Scientific, Cat # 22-363-547) were used to prevent cells from clumping.

### RNA preparation for *in vitro* chemical probing analysis

RNA was natively purified as described before (20,21). Briefly, full-length SChLAP1 RNA was *in vitro* transcribed using recombinant T7 RNA polymerase (in-house) and pBS vector containing SChLAP1 (JX117418.1 base 1-1436). The plasmids containing SChLAP1 fragments were generated by site-directed mutagenesis (New England Biolabs, Cat # E0554S). **Supplementary Table S1** includes the primer sequences used for the mutagenesis.

Plasmids were linearized with the appropriate restriction enzyme (New England Biolabs, Cat # R3136S). The *in vitro* transcription reaction was conducted in 40 mM Tris-HCl (pH 8.0), 12 mM MgCl_2_, 2 mM spermidine, 10 mM NaCl, 10 mM Triton X-100, 10 mM DTT, 3 mM of each NTP, and 20 U of RNase inhibitor (Thermo Fisher Scientific, Cat # 10777019). In-house prepped T7 RNA polymerase (0.1 mg/mL) was added and the reaction was incubated for 2.5 hr at 37°C. After *in vitro* transcription, the sample was treated with DNase (Thermo Fisher Scientific, Cat # AM2238) at 37°C for 30 min. Subsequently, the mixture was supplemented with Proteinase K (Roche Life Science, Cat # 3115836001) for another 30 min at 37°C. The resulting products were filtered (MilliporeSigma, Cat # UFC510008) and SChLAP1 was pooled using size-exclusion chromatography by removing aggregates and prematurely terminated transcripts.

### Chemical probing of *in vitro* transcribed SChLAP1 RNA

For SHAPE, freshly purified RNA was pooled in 25 mM HEPES, pH 7.4, 150 mM KCl, 1 mM EDTA, and 12 mM MgCl_2_ (22,23). RNA was incubated at 37°C for 30 min and divided into two tubes. A final concentration of 200 mM NAI (MilliporeSigma, Cat # 03-310) was added to one tube. An equal amount of pure DMSO (VWR International, Cat # 67-68-5) was added as a control to the remaining tube. DMS probing was conducted as for SHAPE but with 0.4% DMS. Samples were incubated for 10 min at 37°C and retrieved using Zymo RNA Clean & Concentrate Kit (Zymo, Cat #R1015).

### Chemical probing of cellular SChLAP1 RNA

0.8 million cells (per well) were seeded in a 6-well plate 24 hr before chemical probing (24). After washing once with DPBS, cells were resuspended in DPBS supplemented with 80U RNase inhibitor. A final concentration of 0.4% DMS was added to the treated cells, and the same amount of pure EtOH was added to the control cells. Cells were incubated for 6 min at 37°C. Cells were then quickly moved to 4°C, and the reaction was quenched by adding the same volume of quenching solution (1.43 M β-mercaptoethanol) as DMS. Cells were washed once with the quenching solution at 4°C and then incubated in Trizol Reagent (Thermo Fisher Scientific, Cat #15596026) for 5 min at room temperature. Subsequently, total RNA was extracted using the Direct-zol RNA Miniprep kit (Genesee Scientific, Cat # R2050).

### Reverse transcription of modified RNA

For structural probing *in vitro*, reverse transcription was performed using MarathonRT (Kerafast, Cat # EYU007) reverse transcriptase. 300 ng of RNA was mixed with 1 μL of 2 μM gene-specific primers. The mixture was briefly incubated at 95°C for 30 sec and placed immediately on ice.

Next, the RT master mix (50 mM Tris-HCl, pH 8.3, 200 mM KCl, 5 mM DTT, 1 mM MnCl_2_, 10 U Marathon RT, 0.5 mM dNTP, 20% glycerol) was added to a final volume of 10 μL. The samples were incubated at 42°C for 3 hr, and the reverse transcriptase was inactivated by incubation at 70°C for 15 min.

For structural probing *in vivo*, reverse transcription was performed using MarathonRT. 1 μg of total RNA was mixed with 1 μL of 2 μM gene-specific primers. The mixture was incubated for 30 sec at 95°C and placed immediately on ice. Next, 1 μL of 10 mM dNTP, 5 μL 2X RT buffer (100 mM Tris-HCl, pH 7.5, 400 mM KCl, 10 mM DTT, 2 mM MnCl_2_, 40% glycerol), and 10U MarathonRT were added to the final reaction volume of 10 μL. The samples were incubated at 42°C for 3 hr, and the reverse transcriptase was inactivated by incubation at 95°C for 1 min.

After reverse transcription, cDNAs were purified using G-25 spin columns (Cytiva, Cat # 27-5325-01), according to the manufacturer’s instructions.

### Library preparation and high-throughput sequencing

cDNA was PCR-amplified using gene-specific primers and Q5 hot start DNA Polymerase (New England Biolabs, Cat # M0493S). The number of PCR amplification cycles was limited to 25 or less. PCR products were gel-purified, and quantified using Qubit 4 Fluorometer (Thermo Fisher Scientific, Cat # Q33238). 10 ng of cDNA was used for library preparation using Nextera XT DNA Library Preparation Kit (Illumina, Cat # FC-131-1024), according to the manufacturer’s instructions with minor modifications. AMPure XP beads (Beckman Coulter, Cat # A63880) were used for library cleanup. After samples were quality-checked through a bioanalyzer, the NGS library was sequenced at the Drexel Genomics Facility. **Supplementary Table S1** includes all the primers used in this paper.

### SHAPE data analysis

SHAPE reactivity data was processed with the ShapeMapper software (22,23). All replicates passed the ShapeMapper quality control checks. RNA structure prediction and Shannon entropy analysis were performed using SuperFold and SHPAE reactivities as constraints (22,23).

SHAPE differences were determined using the DeltaSHAPE software (25).

### Conservation analysis

The SChLAP1 gene in the human genome (hg38 version) was aligned to and compared with genomes of various vertebrates using the UCSC genome browser (26). Using its table browser tools, SChLAP1 gene coordinates were sent to the GALAXY web-based computational analyses tool, which helped extract the aligned sequences (27). The resulting sequence alignment was converted from MAF to FASTA format. Jalview was used to visualize the conserved sequences (28).

### Plasmid constructs for knockdown and overexpression analysis

For knockdown analysis, Custom-made SChLAP1 targeting shRNA plasmids, along with the non-targeting scramble plasmid, were purchased from GeneCopoeia (Rockville, MD, USA). Following are the target sequences: shSChLAP1-1 (CCAATGATGAGGAGCGGGA), shSChLAP1-2 (CTGGAGATGGTGAACCCAA), and Scramble (GCTTCGCGCCGTAGTCTTA).

For overexpression analysis, the full-length SChLAP1 gene (JX117418.1) was cloned into a lentiviral vector pLenti-GIII-CMV (Applied Biological Materials, Cat # 37847061) to produce pLenti-GIII-FL. pLenti-GIII-CMV backbone was used as a control.

A plasmid containing the 5’ end of SChLAP1 was generated using restriction cloning. The full-length SChLAP1 plasmid was digested with XbaI (New England Biolabs, Cat # R0145S) at 37°C for 1 hour, and bands were separated on a 1% agarose gel. The correct-sized DNA band was cut out and recovered using the Zymoclean Gel DNA Recovery kit (Zymo, Cat # D4001). Eluted DNA was ligated to produce the pLenti-GIII-5’end plasmid.

A plasmid containing the 3’ end of SChLAP1 was by site-directed mutagenesis (New England Biolabs, Cat # E0554S). **Supplementary Table S1** includes the primer sequences used for the mutagenesis.

### shRNA-mediated SChLAP1 knockdown analysis

LNCaP cells were transfected with Lipofectamine 3000 Transfection Reagent (Thermo Fisher Scientific, Cat # L3000008) according to the manufacturer’s instructions. Briefly, 0.7 × 10^6^ cells were seeded in a 6-well plate 24 hr before transfection. 4 μg of shRNA plasmids (hereafter referred to as shSChLAP1) were used for transfection. After 48 hr, the cells were treated with puromycin (2 μg/mL) and incubated further for 48 hr.

### Overexpression analysis

0.7 million cells were seeded in a 6-well plate 24 hr before transfection. LNCaP cells were co-transfected with 1.25 μg of shSChLAP1 and 1.25 μg of an appropriate SChLAP1 overexpression plasmid. For 22Rv1 cells, 2.5 μg of appropriate SChLAP1 overexpression plasmids were transfected. Per the manufacturer’s protocol, the transfection was performed using Lipofectamine 3000 Transfection Reagent (Thermo Fisher Scientific, Cat # L3000001). Puromycin (3 μg/mL) was treated 24 hr post-transfection and cells were incubated for another 48 hr. After GFP signals were checked, cells were collected for downstream analysis.

### Cell proliferation assay

For proliferation assays, cells were collected 72 hr after transfection and plated at 10,000 cells per well density in a 24-well plate. Every 24 hours cells were harvested by trypsinization and counted using a hemocytometer. All experiments were performed in biological triplicate, and cell counts for each biological replicate were the average of technical 2-3 replicates.

### Cell invasion assay

Transwell invasion assay was performed according to the manufacturer’s instruction (Corning, Cat # 354165). Briefly, cells were harvested 72 hr after transfection and stained for 30 min with CellTracker Deep Red dye (Thermo Fisher Scientific, Cat # C34565). 1.2×10^5^ LNCaP cells and 5×10^4^ 22Rv1 cells in serum-free DMEM were added to the Matrigel-coated upper chambers. As a chemoattractant, DMEM containing 5% FBS was added to the lower chamber. After 20 hr, the invaded cells were imaged using an EVOS FL Auto microscope (Life Technologies). The invasive abilities were evaluated by counting the number of invading cells from three random fields per transwell. Experiments were performed in a biological quadruplicate.

### RNA extraction and RT-qPCR

Cells transfected with appropriate plasmids were trypsinized and pelleted. Total RNA was extracted using the Direct-zol RNA Miniprep kit (Genesee Scientific, Cat # R2050) following the manufacturer’s protocol. The quality and concentration of RNA were determined using a NanoDrop One/OneC Microvolume UV-Vis Spectrophotometer (Thermo Fisher Scientific).

Reverse transcription was performed using iScript Advanced cDNA Synthesis Kit (BioRad, Cat # 1725037). The resulting cDNA was used to perform qPCR analysis using SsoAdvanced Universal SYBR Green Supermix (BioRad, Cat #1725270) and a Bio-Rad CFX-96 real-time PCR detection system. The expression levels of the target genes were normalized to the transcription levels of GAPDH. The 2^−ΔΔCt^ method was used to calculate the relative gene expression levels. **Supplementary Table S1** includes all the primers used in this paper.

### Statistical analysis

Statistical analysis was performed using GraphPad Prism 10.3.1 software (GraphPad Software, Inc.). Student’s *t*-test was used to compare the means of biological triplicate or quadruplicate, with each replicate a representative mean of 2-3 technical replicates. Data are represented as the mean ± standard error of the mean (SEM). Statistical significance was defined as follows: \**p*≤0.05, \*\**p*≤0.01, \*\*\**p*≤0.001, \*\*\*\**p*≤0.0001, with “ns” indicating no significance. Where statistical significance is not identified, data is assumed to have a *p*-value greater than 0.05.

## RESULTS

### SChLAP1 is purified as a homogeneous population under non-denaturing conditions

To determine the structure of large RNAs, the first challenge is to purify homogeneous, monodisperse samples while preserving their native, functional structures. Since the traditional denaturation-refolding purification methods can cause misfolding and aggregation, non-denaturing methods have been developed, such as affinity tag-based as well as native purifications (20,29,30). The native purification method was successfully applied for a handful of lncRNAs (20,21). As such, we used the native purification method to purify SChLAP1 (**Figure 1A**). After *in vitro* transcription, SChLAP1 was purified by size-exclusion chromatography (SEC) by pooling the fractions containing the RNA of interest (**Figure 1B**).

**Figure 1.**
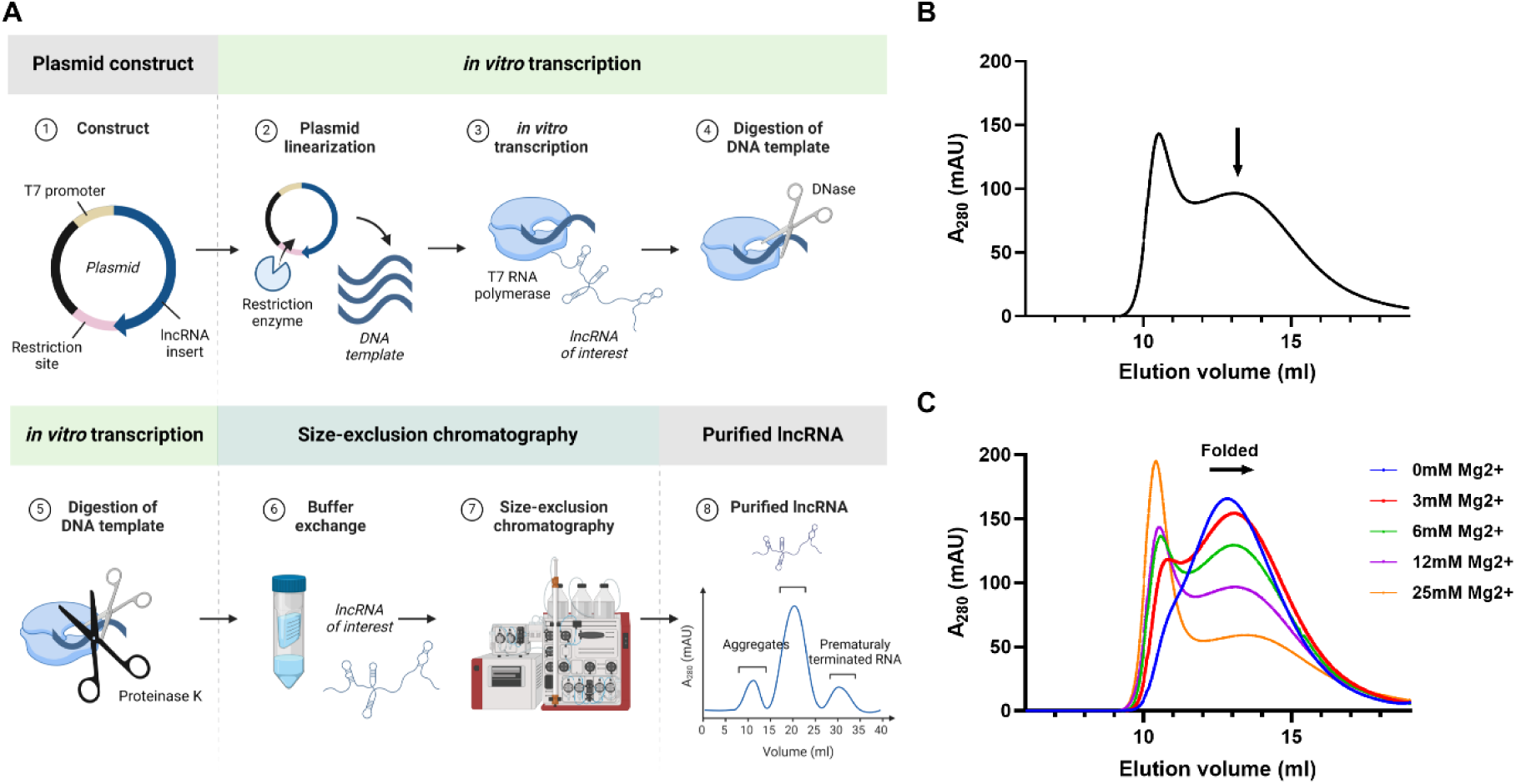
Purification of the lncRNA SChLAP1. (A) Overview of SChLAP1’s native purification method. (B) SChLAP1 was purified by size-exclusion chromatography (SEC) following *in vitro* transcription. (C) SEC profiles of SChLAP1 folded at varying concentrations of Mg^2+^. Created with BioRender.com.

Divalent cations, such as Mg^2+^, are essential for the folding and stability of RNA molecules, we determined that SChLAP1 could be folded at various Mg^2+^ concentrations (31,32). While the homogeneity of RNA was consistent across a wide range of Mg^2+^ concentrations, higher concentrations of Mg^2+^ caused a slight rightward shift of chromatograms, with an increase in aggregation and a decrease in absorbance (**Figure 1C**). This indicates that SChLAP1 folded into more compact structures through intramolecular interactions, suggesting that SChLAP1 may have more complex, higher-order structures (33). Taken together, we successfully purified SChLAP1 using the native purification protocol.

### Chemical probing of SChLAP1

For secondary-structure determination of SChLAP1, we used chemical probing followed by mutational profiling (MaP), with DMS (dimethyl sulfate) and NAI (2-methylnicotinic acid imidazolide; SHAPE reagent) as RNA-modifying chemicals (22,34). These probes modify the single-stranded regions of RNA, inducing reverse transcriptase enzymes to misread and incorporate random mutations into the newly synthesized cDNAs. After next-generation sequencing, the resulting mutations are counted by mapping the reads to the original RNA sequence, thereby generating reactivities. High SHAPE reactivity refers to flexible, single-stranded structures, whereas low SHAPE reactivity refers to double-stranded structures (22,23). Our results indicate that SChLAP1 was successfully chemically modified, as demonstrated by higher mutation rates in the probe sample compared to the control (**Figure 2A**). Further, SHAPE and DMS reactivities from independent experiments are highly reproducible, with Pearson correlation coefficients (*r*) of 0.95 and 0.99, respectively (**Figure 2B and 2C**).

**Figure 2.**
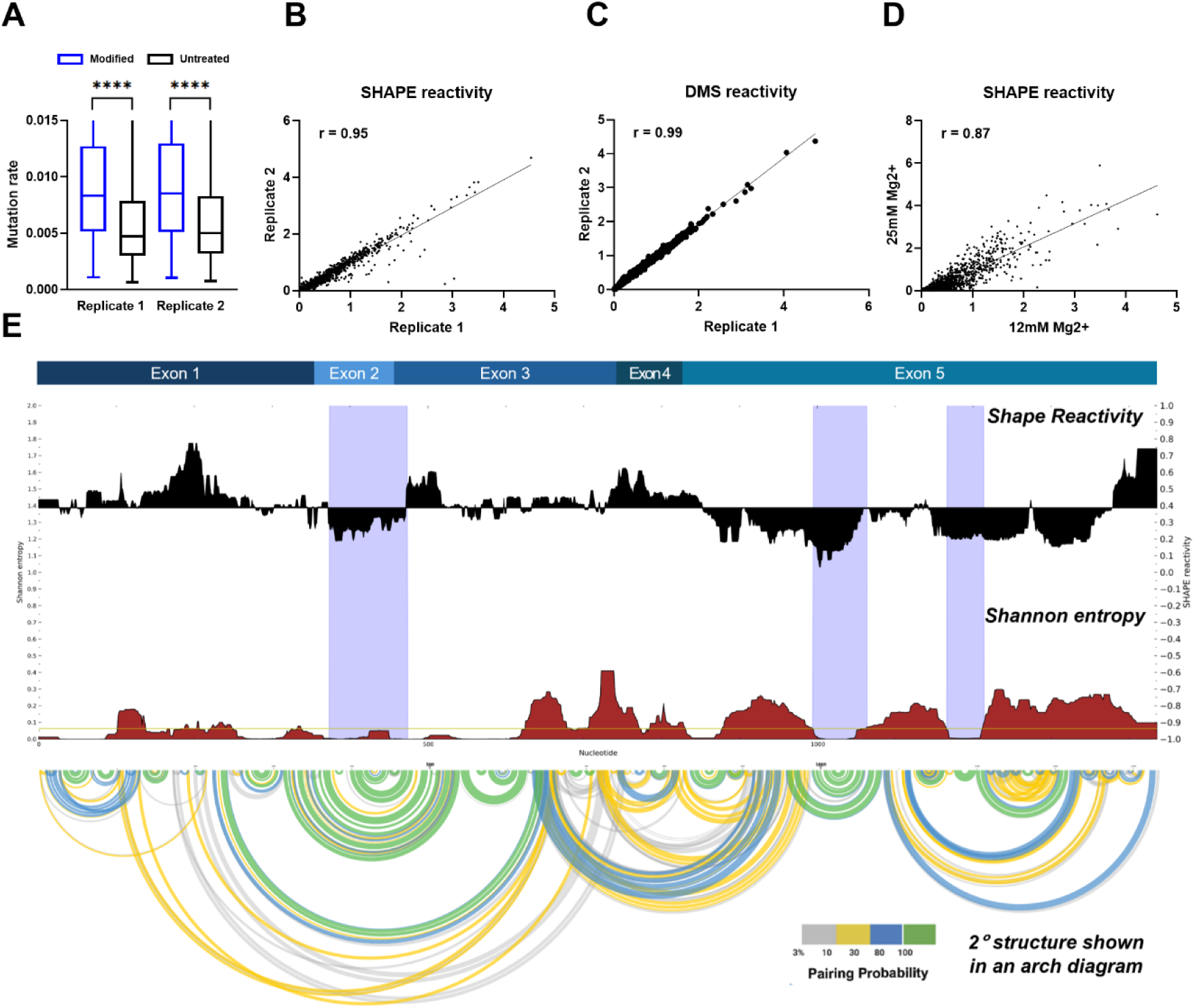
*in vitro* structural probing of SChLAP1 using SHAPE-MaP analysis. (A) SChLAP1 was chemically modified successfully. As expected, modified samples show higher mutation rates compared to control samples. (B) SHAPE reactivities were reproducible among different biological replicates. Pearson correlation coefficient (*r*) values are indicated. (C) DMS reactivities were reproducible among different biological replicates. Pearson correlation coefficient (*r*) values are indicated. (D) SHAPE reactivities at 12 mM and 25 mM Mg^2+^ are strongly correlated (*r* = 0.87). (E) SChLAP1 has three highly structured and stable areas with low SHAPE reactivities and low Shannon entropies (violet-shading).

A comparison of SHAPE reactivities for SChLAP1 under two Mg^2+^ concentrations (12 mM and 25 mM) in the folding buffer revealed a strong correlation (Pearson *r* = 0.87), suggesting that the secondary structure is preserved even at higher [Mg^2+^] (**Figure 2D**). Additionally, chemical probing of SChLAP1 in the higher [Mg2+] folding buffer showed strong agreement in both SHAPE and DMS reactivities across different biological replicates (**Supplementary Figure S1**).

### SChLAP1 exhibits an intricate secondary structure with unique sequential characteristics at the 5’ end

Next, SHAPE reactivities were used as constraints for secondary structure prediction using SuperFold (**Figures 2 and 3**) (23). Our *in vitro* secondary structural map reveals that SChLAP1 is a highly structured lncRNA, with 49.7% of its nucleotides base-paired (**Figure 3**). Notably, SChLAP1 features three highly structured and stable regions with low SHAPE reactivities and low Shannon entropies (violet shading; see **Figure 2E**).

**Figure 3.**
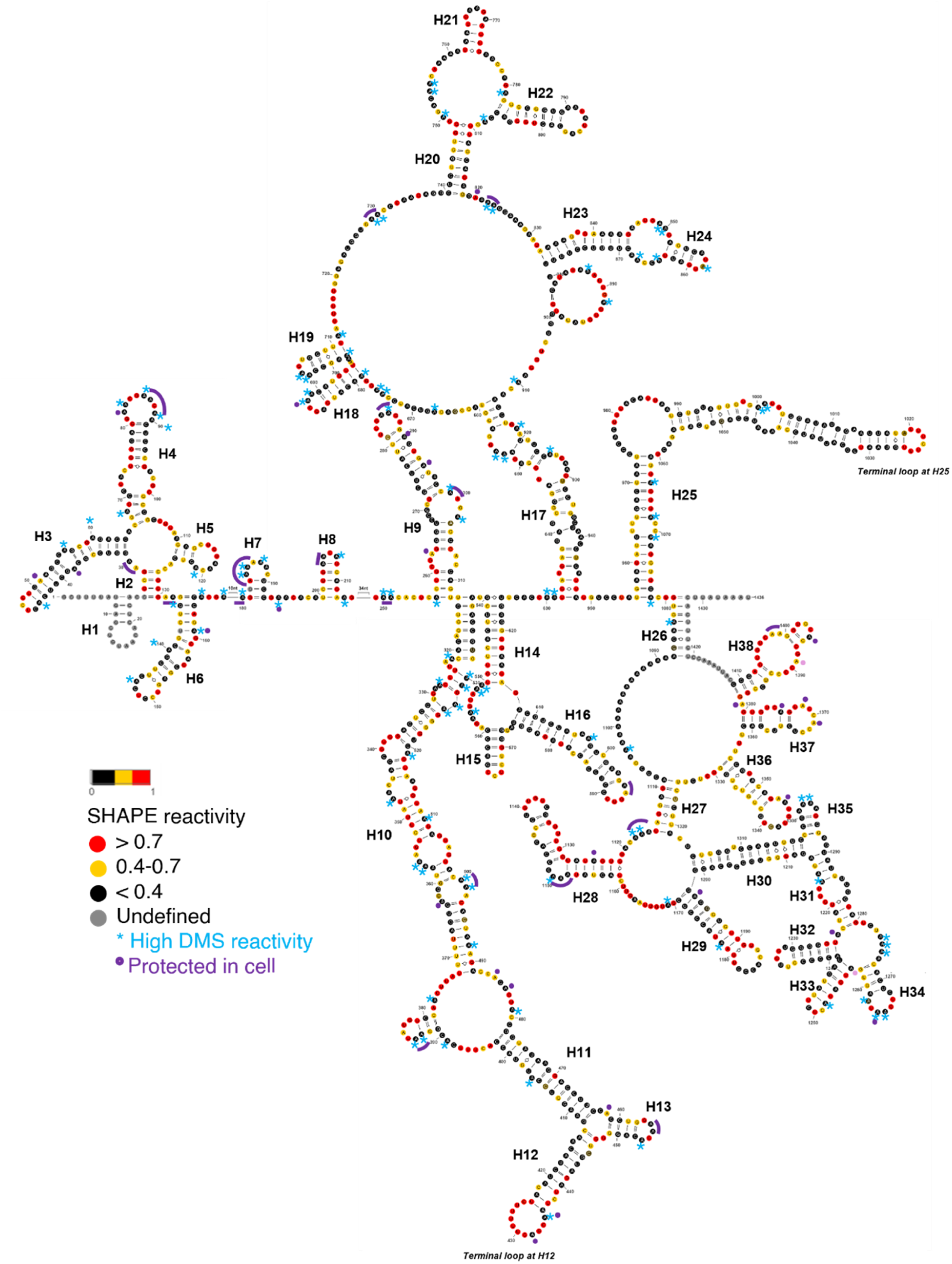
*in vitro* secondary-structure model of SChLAP1 (averaged of two biological replicates). Nucleotides with high SHAPE reactivity are highlighted in red, nucleotides with medium SHAPE reactivity are highlighted in yellow, nucleotides with low SHAPE reactivity are highlighted in black, and nucleotides with ‘no data’ are highlighted in grey. The asterisk represents nucleotides with low SHAPE reactivity and high DMS reactivity. Nucleotides protected during *in vivo* DMS probing experiments are highlighted in purple.

Along with its intricate structure, SChLAP1 exhibits unique sequential characteristics at its 5’ end, including repeats and conservation. Notably, the first exon of SChLAP1 comprises three large repeat sequences (**Supplementary Figures S2A through S2D**). LncRNA repeats, as observed in the well-studied lncRNA Xist, have been shown to act as possible lncRNA-protein interaction hubs, thereby contributing to the overall functionality of lncRNA (35,36).

Moreover, the 5’ end of SChLAP1 has evolutionarily conserved regions; specifically, the second exon is conserved in 57 out of 100 vertebrates, making it the most conserved among the five exons of SChLAP1 (**Supplementary Figure S2E**). Furthermore, we found multiple highly conserved motifs throughout the second exon (**Supplementary Figure S2F**). We also noted that SChLAP1 was conserved only among mammals, not in other vertebrates such as Zebrafish.

### SChLAP1 DMS reactivities are in good agreement with the directed secondary structure model

DMS reactivities were also in good agreement with the SHAPE reactivity of SChLAP1. No nucleotides (A) with high SHAPE reactivity (greater than 0.7) exhibited low DMS reactivity (less than 0.4). However, 28.7% of nucleotides (A) with low SHAPE reactivity showed high reactivity to DMS. This may be because SHAPE and DMS target different chemical sites and aspects of RNA structure. DMS focuses on base accessibility, particularly for A, while SHAPE measures the flexibility and dynamics of the ribose backbone. Therefore, the combination of lower SHAPE reactivity and higher DMS reactivity suggests that these adenines may be unpaired or single-stranded but structurally constrained by nearby interactions or tertiary structure elements, reflecting the structural environment of SChLAP1. Indeed, all these A’s with high DMS reactivity are located within loop regions or at the beginning or end of the loop regions, supporting our secondary structural model (blue asterisk; see **Figure 3**). In summary, SHAPE-MaP analyses were used to determine the secondary structure of SChLAP1, and DMS probing results were consistent with the SHAPE-directed secondary structure model.

### *In vivo* chemical probing reveals protein-binding hotspots on SChLAP1

While *in vitro* structure probing is useful for mapping the thermostable state of RNA determined mainly by its primary sequence, *in vivo* structure probing enables the characterization of biologically relevant RNA conformations (24). This is because the presence of RNA chaperones as well as interacting partners, such as proteins, DNAs, and other RNAs, contributes to the RNA folding in cells (37). Therefore, we probed SChLAP1 *in vivo* using DMS and compared the *in vitro* and *in vivo* DMS reactivities to identify potential protein-binding hubs (**Figure 4A**). By subtracting *in vivo* DMS reactivities from *in vitro* DMS reactivities, we calculated the delta (Δ) DMS reactivity. Positive Δ values indicate the nucleotides are more protected in cells, suggesting protein-binding, whereas negative Δ values indicate greater flexibility in cells.

**Figure 4.**
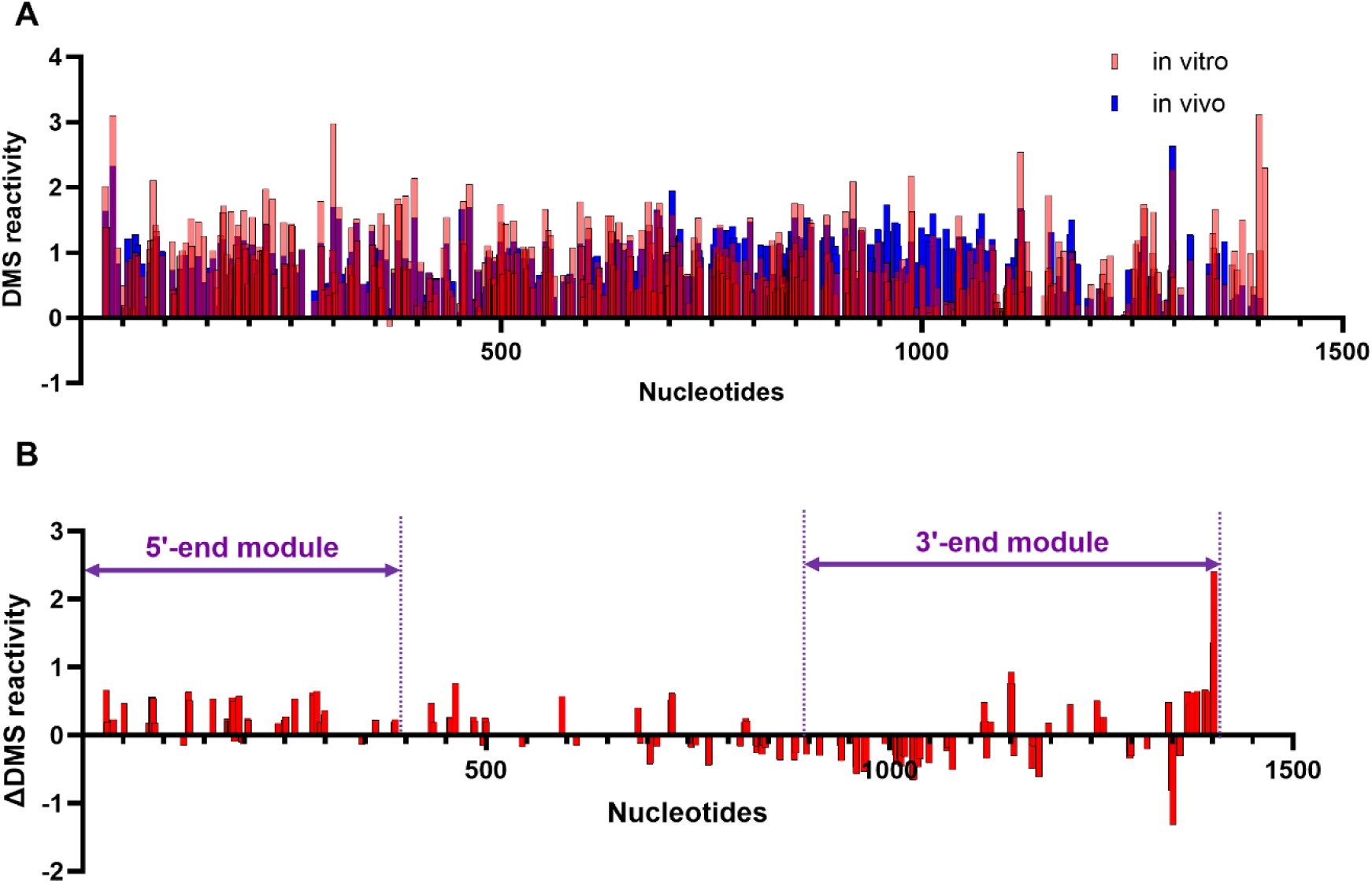
*in vivo* chemical probing reveals protein-binding hotspots on SChLAP1. (A) Comparison of *in vitro* and *in vivo* DMS reactivities. (B) The 5’ and 3’ ends of SChLAP1 showed protection, suggesting potential protein-binding sites within these regions.

Our findings reveal protein hotspots within the two terminal modules at the 5’ and 3’ ends of SChLAP1 (**Figure 4B**). These potential protein-binding sites are also highlighted in violet in **Figure 3**. It is important to note that our results specifically target A bases, meaning nearby regions may also be involved in protein interactions.

Among the 429 A bases in SChLAP1 (excluding primer-binding sites), only 138 (32%) showed significant differences in DMS reactivity between *in vitro* and *in vivo* conditions. Of these, 70 bases were protected in cells, while 98 were more reactive. The regions identified include the terminal loops of the two terminal modules. Notably, we identified potential SChLAP1-protein binding sites spanning the first exon (Nucleotides 1-338) (**Figure 4B**). The conserved second exon (Nucleotides 339-433), particularly the terminal loop at the helix 12 (Nucleotides 424-436), was also protected in cells (**Figure 3 and Figure 4B**). Altogether, our *in vitro* probing studies identified structurally stable regions at the 5’ and 3’ ends of SChLAP1, and our *in vivo* probing studies reveal that some of these regions are protected in cells, suggesting they may be potential protein-binding sites.

### The 5’- and 3’-end modules of SChLAP1 have independently folding structural elements with potential functional roles

To further investigate whether these terminal modules have independently folding domains, we designed fragments encompassing the secondary structure elements of these regions and performed SHAPE analysis. Specifically, we made two fragments, Fragment 1 (Nucleotides 222-651) and Fragment 2 (Nucleotides 956-1428). Both fragments were transcribed and folded alongside the full-length transcript (**Figures 5A and 5D**) (38).

**Figure 5.**
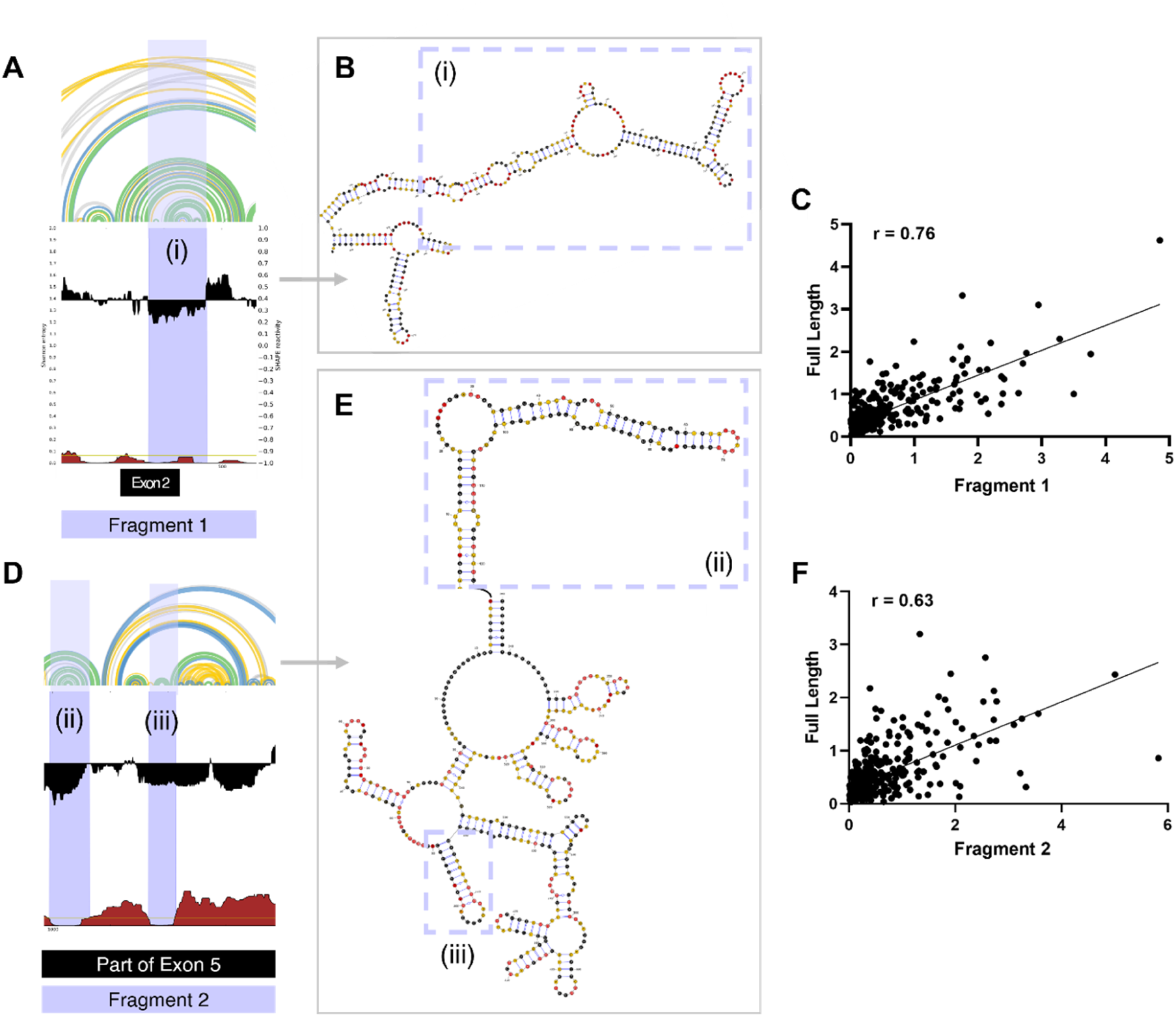
SHAPE analysis of SChLAP1 fragments. (A) SHAPE reactivities and Shannon entropies for Fragment 1 (Nucleotides 222-651). (B) The secondary structure of Fragment 1 reveals distinct structures, characterized by a multi-branched loop with stems and internal loops. (C) A scatter plot comparing SHAPE reactivities of Fragment 1 with the corresponding region in full-length SChLAP1. Pearson correlation coefficient (*r*) values are indicated. (D) SHAPE reactivities and Shannon entropies for Fragment 2 (Nucleotides 956-1428). (E) The secondary structure of Fragment 2 reveals distinct structures with stem-loop motifs. (F) A scatter plot comparing SHAPE reactivities of Fragment 2 with the corresponding regions in full-length SChLAP1. Pearson correlation coefficient (*r*) values are provided.

Comparison of SHAPE reactivities for Fragment 1 with the corresponding region in the full-length SChLAP1 revealed a strong correlation (Pearson *r* = 0.76) (**Figure 5C**). Particularly, the region with low reactivities and low entropies exhibited a higher correlation (Pearson *r* = 0.81), forming a structurally stable region with a multi-branched loop with stems and internal loops (**Supplementary Figure S3A and Figure 5B**).

Similarly, SHAPE reactivities for Fragment 2 also showed strong correlations with their corresponding regions in the full-length SChLAP1 (Pearson *r* = 0.63). (**Figure 5F**). Focusing on the regions with low Shannon entropy, they exhibited notably higher correlations, with Pearson *r* values of 0.75 and 0.85, respectively (**Supplementary Figures S3B and S3C**). This further supports the conclusion that Fragment 2 has independently folded, structurally stable regions forming two stem-loop motifs (ii and iii; see **Figure 5E**). In conclusion, the terminal modules exhibit a mix of structured and unstructured regions, with the interplay between the two likely holding functional significance.

### Structural and functional characterization of the structured regions in SChLAP1

#### The 5’-end module

Next, we set out to test the functional significance of these structured regions identified from our probing experiments. First, we investigated whether the three repetitive regions in exon 1 of the 5’ module form the same secondary structure. While the first two repeats appear to adopt a similar structure, the last repeat is predicted to fold differently, with an elongated stem-loop structure (56 nucleotides from 257 through 312 bases; see **Supplementary Figure S4A**). This could be due to two reasons: first, the primer-binding site is located near the start of the first repeat, which may have prevented us from obtaining reliable reactivity data. Second, the high Shannon entropies around the repeat regions indicate that these regions possess dynamic structures.

Another intriguing structure is the aforementioned second exon region. Unlike small, autonomous helical stems (e.g. H7 and H8; see **Figure 3**), some larger structures are formed through long-range base-pairing. The final 26 nucleotides of the first exon and the entirety of the second exon participate in long-range interactions with nucleotides 538 to 436 (part of the third exon, in reverse order), resulting in the formation of intricate structures comprising 4 helices, 3 terminal loops, and 2 three-way junctions (**Figure 3 and Figure 5B**). Notably, the terminal loop at H12 within this structure is a U-rich region (**Figure 3**). U-rich sequences are known to serve as recognition motifs for various RNA-binding proteins (RBPs), which regulate different aspects of RNA biology, including splicing, export, localization, stability, and translation (39,40). This loop is also identified as one of the potential protein-binding sites based on our *in vivo* chemical probing experiment (**Figure 4**).

In our 5’ module, 201 out of 433 nucleotides were base-paired (46.4%). Overall, based on its sequential and structural characteristics, we hypothesized that the first and second exons (hereafter referred to as 5’ end) are an important structural module of SChLAP1 with a potential functional significance.

#### The 3’-end module

The structural module at the 3’ end of SChLAP1 is characterized by relatively low SHAPE reactivities and a mix of low and high Shannon entropy. These findings indicate that the 3′-end region has structured regions surrounded by flexible and dynamic domains. The two structured stem-loop elements are highlighted (**Figure 5D**). Additionally, this region (exon 5) contains several conserved elements (**Supplementary Figure S2E**). Our *in vivo* chemical probing results also identified another protein-binding hub within the 3’ end, though less prominent than the 5’ end (**Figure 4**).

Specifically, the structure formed at the start of this module (area (ii); see **Figure 5E**) also merits consideration. The terminal loop at H25 (Nucleotides 1019-1026) in this region is U-rich (**Figure 3**). However, the potential protein-binding sites in this terminal loop at the 3’ end were not detected in our *in vivo* DMS probing results, as this region lacks A bases (**Figure 4**). Further research into this structure could therefore provide valuable insights into the functional importance of this structure.

Stretches of adenosine (A) are present in the internal loop at the 3’ end (Nucleotides 1088-1104; see **Figure 3**). Interestingly, these regions exhibit low reactivity to chemical reagents yet appear to be part of huge single-stranded loops. This could suggest the presence of a complex tertiary structure that is difficult to detect through chemical probing. Functionally, these A stretches may play important roles in processes such as RNA stability and decay. For example, A-rich sequences are known to regulate viral RNA replication or interact with host cell machinery to promote translation and viral gene expression (41). Together, these findings led us to hypothesize that SChLAP1 consists of two distinct structural modules, the 5’ and 3’ ends, each featuring significant secondary structural content with well-defined motifs that may play critical functional roles.

### The 5’ and 3’ ends of SChLAP1 promote prostate cancer cell invasion

Previous research revealed that SChLAP1 could modulate prostate cancer cell invasion and proliferation (19). Therefore, to examine the potential functional significance of the 5’ and 3’ ends of SChLAP1 in prostate cancer, we overexpressed plasmids containing these regions in parallel with full-length in two cell lines, LNCaP and 22Rv1. An empty vector was used as a control. We chose the 22Rv1 cell line, a human prostate carcinoma epithelial cell line with low endogenous SChLAP1 expression, making it suitable for overexpression, and LNCaP, an androgen-sensitive prostate adenocarcinoma cell line, which naturally exhibits high SChLAP1 expression (**Figure 6A**). As such, to minimize the potential confounding effects of basal expression of SChLAP1 in LNCaP cells, we first knocked down SChLAP1 and then overexpressed either full-length SChLAP1 or each of its ends (**Figure 6B and 6C**). The overexpression efficiency was verified by RT-qPCR analysis 72 hr post-transfection (**Figure 6C and 6D**).

**Figure 6.**
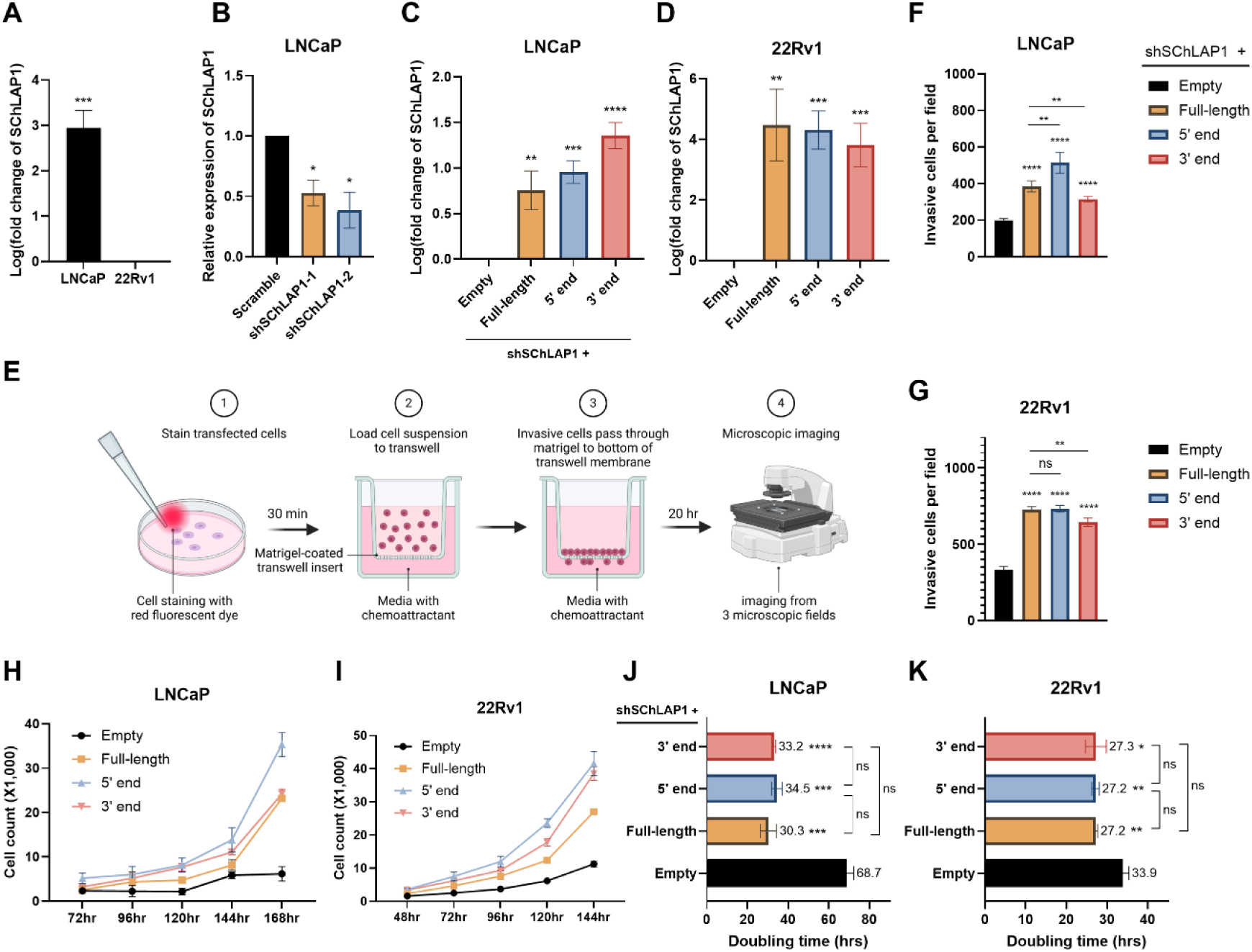
The 5’ and 3’ ends of SChLAP1 independently promote prostate cancer cell invasion and proliferation. (A) SChLAP1 expression levels in LNCaP and 22Rv1 cells were measured by RT-qPCR analysis. (B) The efficiency of shRNA-mediated SChLAP1 knockdown in LNCaP cells was evaluated via RT-qPCR. (C) The co-transfection efficiency of SChLAP1 knockdown and overexpression in LNCaP cells was assessed via RT-qPCR. (D) The overexpression efficiency of SChLAP1 in 22Rv1 cells was assessed via RT-qPCR. (E) Overview of transwell cell invasion assays. (F & G) Cell invasion upon overexpression of full-length SChLAP1 or fragments was assessed using transwell invasion assays in both LNCaP (F) and 22Rv1 (G) cells. (H & I) Cell proliferation was evaluated using cell counting analysis in LNCaP cells (H) and 22Rv1 cells (I) overexpressing either full-length SChLAP1 or its fragments, with an empty vector used as the control. See Supplementary Figure S6 for bar plots depicting these data. (J) Doubling time of LNCaP cells upon overexpressing the 5’ or 3’ end of SChLAP1. (K) Doubling time of 22Rv1 cells upon overexpressing the 5’ or 3’ end of SChLAP1. Data are represented as mean ± SEM from three biological replicates; \**p*≤0.05, \*\**p*≤0.01, \*\*\**p*≤0.001, \*\*\*\**p*≤0.0001, with “ns” indicating no significance.

First, we performed transwell invasion assays to analyze the effects of the 5’ and 3’ ends of SChLAP1 on cell invasion (**Figure 6E**). We noticed cell invasion was increased by both the 5’ and 3’ end of SChLAP1 overexpression, compared to the control group (**Figures 6F, 6G, and Supplementary Figures S5**). Interestingly, in LNCaP cells, the 5’ end exhibited a significantly higher effect on cell invasion than the full-length, while the 3’ end showed a markedly smaller effect than the full-length. In 22Rv1 cells, the effects of the 5’ end and the full-length were comparable, whereas the 3’ end had a significantly reduced impact compared to both (**Figure 6F and 6G**). These data demonstrate that the 5’ and 3’ ends of SChLAP1 can promote prostate cancer cell invasion, with the 5’ end playing a particularly significant role.

### The 5’ and 3’ ends of SChLAP1 promote prostate cancer cell proliferation

Next, we determined the effect of the 5’ and 3’ ends of SChLAP1 on cell proliferation using direct cell counting and doubling rate analysis. These results revealed that overexpressing either end of SChLAP1 significantly promoted cell proliferation in both cell lines compared to the control group (**Figures 6H and 6I, and Supplementary Figure S6**). The effects of the ends were comparable to those of the full-length SChLAP1 (**Figures 6J and 6K**). Overall, these data show that both structural modules of SChLAP1 could independently promote prostate cancer cell proliferation.

### The 5’ and 3’ ends of SChLAP1 elevate invasive and proliferative gene expression

To further investigate whether the 5’ and 3’ ends of SChLAP1 regulate the expression of genes involved in prostate cancer cell invasion and proliferation, we examined the expression level of pro-invasion and pro-proliferation genes in LNCaP and 22Rv1 cells upon overexpressing the ends of SChLAP1. To mitigate the impact of increased basal SChLAP1 expression in LNCaP cells, shRNA-mediated knockdown of SChLAP1 was performed prior to overexpressing either end in these cells. Consistent with previous research, our RT-qPCR results demonstrated that SChLAP1 knockdown significantly inhibited the expression of invasion markers MMP-9 and MMP-14, as well as the proliferation marker VEGF, in prostate cancer cells (**Figure 7A**) (42). MMP-9 (matrix metalloproteinase-9) and MMP-14 (matrix metalloproteinase-14) drive tumor invasion and growth by degrading extracellular matrix (ECM) proteins (43,44). VEGF (vascular endothelial growth factor) stimulates angiogenesis by interacting with MMP-9 and MMP-14, promoting tumor metastasis (44).

**Figure 7.**
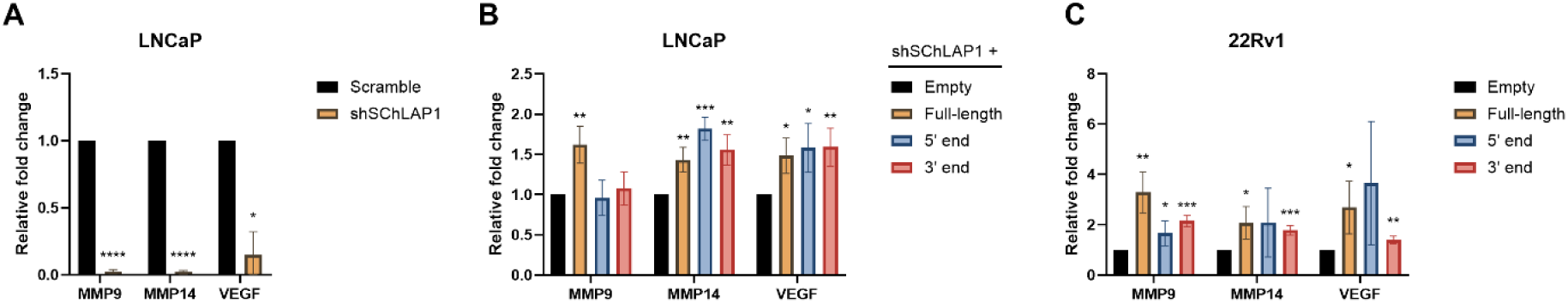
Overexpression of the 5’ and 3’ ends of SChLAP1 increased the expression of invasive and proliferative gene markers in prostate cancer cells. (A) SChLAP1 knockdown significantly reduced the expression of pro-invasion and pro-proliferation genes in LNCaP cells. (B and C) Invasive and proliferative gene expression in LNCaP and 22Rv1 cells was evaluated via RT-qPCR after overexpressing the 5’ or 3’ end of SChLAP1. Data are represented as mean ± SEM from three or four biological replicates; \**p*≤0.05, \*\**p*≤0.01, \*\*\**p*≤0.001, \*\*\*\**p*≤0.0001, with “ns” indicating no significance.

Our RT-qPCR analysis revealed that overexpression of either end of SChLAP1 led to a notable upregulation of the invasion and proliferation markers in both cell lines, although the magnitude of their effects was generally lower than that observed with the full-length (**Figure 7B and 7C**). This data indicates that the 5’ and 3’ ends of SChLAP1 may promote cancer cell invasion by upregulating the expression of invasion and proliferation marker genes.

## DISCUSSION

### The structure-function relationship of lncRNA in prostate cancer

A growing body of scientific evidence emphasizes the significance of lncRNA’s modular structural architecture (16–18). Although the structural studies of lncRNAs are fundamental for their detailed biological mechanisms, the structure-function relationship of lncRNAs has been understudied. Among the approximately 20,000 annotated lncRNAs in the human genome, less than 20 lncRNAs have been investigated using structural approaches (9,36). SChLAP1 is one of the structurally understudied lncRNAs. SChLAP1 is markedly overexpressed in aggressive prostate cancer and has been implicated in promoting the metastatic progression of the disease (19). Since the specific role of SChLAP1 in driving prostate cancer metastasis remains largely unexplored, this study aimed to investigate its structure-function relationship to uncover the underlying mechanism.

### The secondary structural map of SChLAP1 revealed two distinct structural modules

Our *in vitro* structure model revealed that SChLAP1 is a highly structured lncRNA with nearly 50 percent of its nucleotides being base-paired. In detail, SChLAP1 has two distinct structural modules, including 5’ and 3’ ends. Both terminal modules have high secondary-structural content. In addition to their structural elements, the 5’ end module of SChLAP1 is composed of extremely repetitive sequences and has short sequence motifs highly conserved in mammals. Our *in vivo* chemical probing experiments uncovered extensive protein-binding sites at the 5’ and 3’ ends of SChLAP1, further supporting their potential functional significance.

Very recently, the Hargrove lab published a secondary structural model of SChLAP1 in bioRxiv (45). Our model agrees with their model, especially the stable regions with low SHAPE reactivity and low Shannon entropy, which form similar structures in both models (45). However, some differences exist, such as the predicted structure of the repeat regions in the 5’ end. That said, both studies observed high Shannon entropies around repeat regions at the 5’ end, suggesting dynamic structures that may facilitate protein-binding function (45). It is worth pointing out that the terminal loop at H12 in the 5′-end module was identified as a potential protein-binding motif in both studies (45). However, the terminal loop at H25 in the 3′-end module was undetectable by our *in vivo* DMS probing due to the lack of an A-base but was identified using *in vivo* SHAPE (45). The robust structural elements remain consistent despite studies conducted in two different labs and differences in conditions, such as reverse transcriptase and the number of amplicons used for SHAPE-MaP analysis. These findings provide a robust structural model of SChLAP1 and underscore its biological significance.

### SChLAP1’s oncogenicity is mediated by its distinct structural modules

Indeed, our functional studies showed that the truncated versions of SChLAP1 contain all the elements necessary to promote the invasion and proliferation of prostate cancer cells.

Interestingly, the 5’ end of SChLAP1 often exhibits effects comparable to the full-length RNA, whereas the 3’ end shows a smaller functional effect. This trend is particularly evident in the cell invasion assays. Further studies, such as RNA sequencing to investigate changes in relevant pathways, would be valuable to elucidate the reasons behind these differences. Since SChLAP1 is remarkably enriched in the nucleus of the cells (19), we assume that SChLAP1 may be engaged in the transcriptional modulation of upstream regulators of the proliferative and invasive phenotype of cancer cells.

Moreover, in our study, to test the functional significance we employed two cell lines, 22Rv1 cells, a prostate cancer cell line with low basal SChLAP1 expression, and LNCaP cells, which have high basal SChLAP1 expression. These approaches allow us to ensure that our findings are biologically more relevant within a cancer-like microenvironment and not cell-line specific.

### The structural complexity of SChLAP1: balancing organized and flexible modules in protein interactions

Unlike the compact and highly organized structure of lncRNA MUNC, SChLAP1 consists of not only highly organized modules but also long, unstructured domains and repetitive sequences like Xist (18,46). The long, unstructured regions of lncRNA may serve as protein-landing pads while the structured regions could act as scaffolds or offer protein-binding structural motifs, playing crucial roles in important biological pathways.

Both terminal modules exhibit high secondary-structural content, while the intervening regions, referred to as the central module, possess low secondary-structural content. The central module, primarily spanning exons 3 and 4, displays high SHAPE reactivities indicative of low secondary-structure content and corresponding high Shannon entropy, reflecting greater structural flexibility. This module contains short stretches of base-paired regions, forming a large internal loop (Nucleotides 660-913) (**Figure 3**). While many of the identified protein-binding sites lie within the highly structured and well-folded regions of SChLAP1’s terminal modules, the central module contains a few protein-binding sites situated in regions that are neither well-folded nor stable. (**Figure 4**). Such regions may be important for the localization of SChLAP1.

For example, the repetitive elements of the lncRNA Xist, with their dynamic structures, act as adaptable protein-docking sites, promoting protein multimerization to support RNA processing and localization (46).

### Towards tertiary structural studies and interactome analysis

Our secondary structural studies suggest that SChLAP1 may exhibit conformational dynamics, with its complex tertiary structure playing a crucial functional role. While tertiary lncRNA structures remain underexplored, techniques like cryo-EM are aiding in their elucidation (47). The secondary structured model determined in this study provides a valuable map for designing constructs for 3D structural studies.

Equally important is analyzing SChLAP1’s interactome, as protein interactions can stabilize lncRNA conformations and facilitate structural studies. For instance, m6A modifications destabilize RNA duplexes, exposing protein-binding sites that proteins like HNRNPC can stabilize the structures, enabling detailed analysis (48,49). Structural studies of SChLAP1 complexes with binding partners are key to understanding its mechanisms.

### Summary

In summary, our study demonstrates that the proliferative and invasive properties of SChLAP1 are driven by its modular architecture, particularly its terminal modules, which may represent potential targets for small-molecule-based cancer therapies. Ultimately, this work has significant public health implications in that SChLAP1 could be a novel target for prostate cancer therapeutics.

## ACKNOWLEDGMENTS

We thank Dr. Mauricio Reginato and Dr. Alessandro Fatatis at Drexel University College of Medicine for the reagents.

## CONTRIBUTIONS

M.O. and S.S. contributed to concept design and manuscript preparation. M.O., R.K., and Z.C. performed the experiments. S.S. and M.O. performed data analysis.

---

**Supplementary Figure S1.**
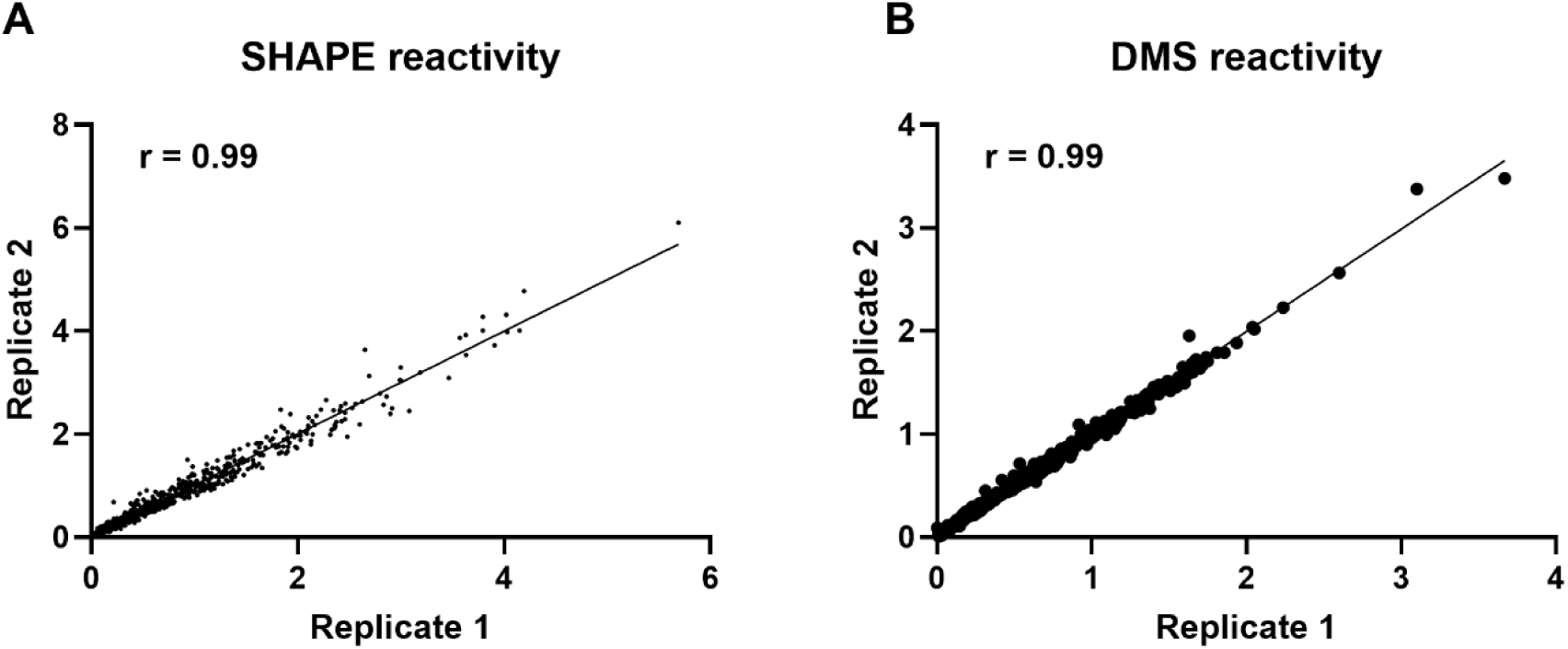
Chemical probing of SChLAP1 in 25 mM Mg^2+^ folding buffer. SHAPE reactivity (A) and DMS reactivity (B) results were reproducible among different biological replicates. Pearson correlation coefficient (*r*) values are indicated.

**Supplementary Figure S2.**
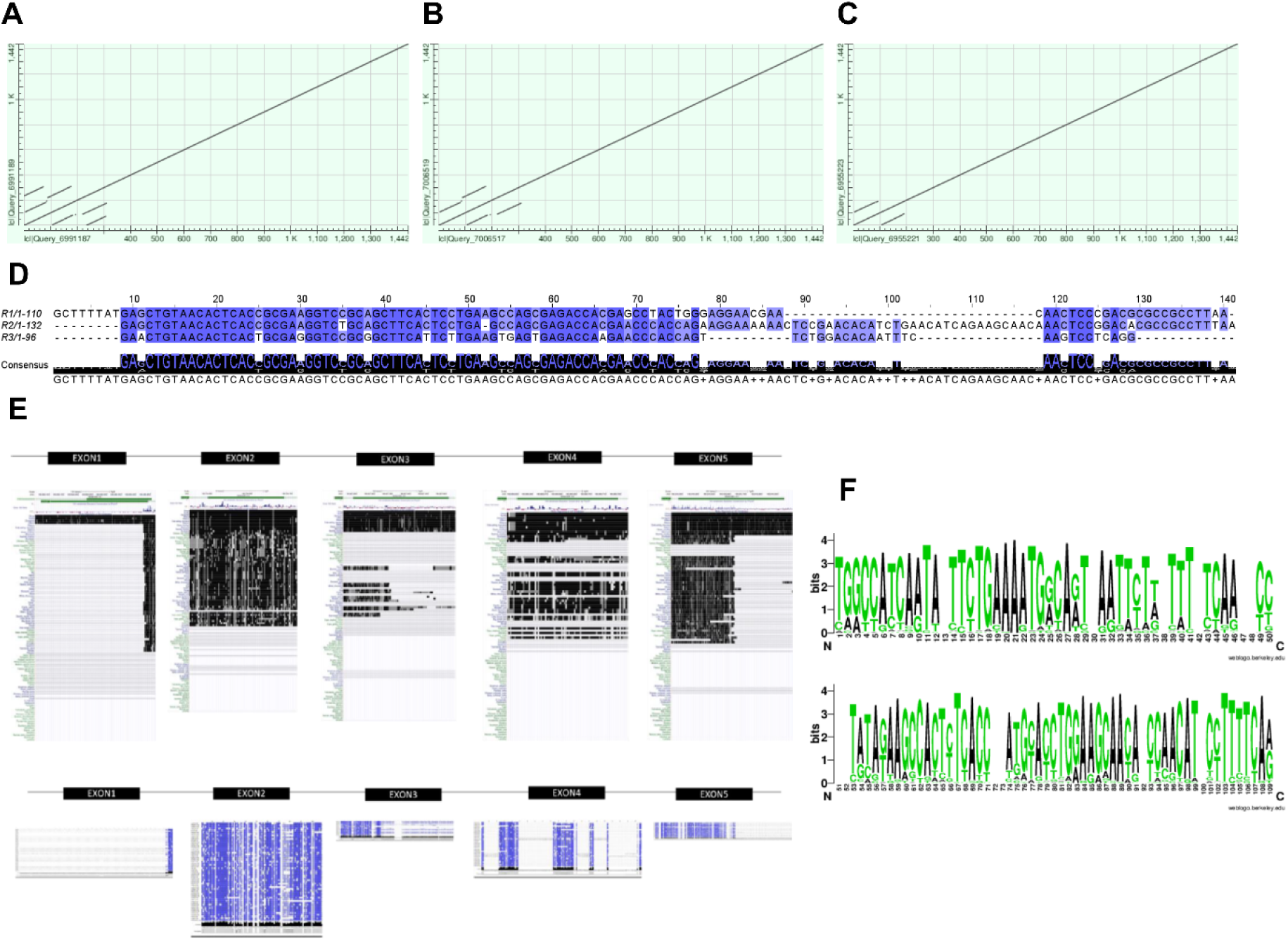
Dot plots of BLAST self-alignment of SChLAP1 sequences with a setting of “somewhat similar (A),” “more dissimilar (B),” and “highly similar (C)”. Comparison of the SChLAP1 sequence with itself revealed the presence of several repetitive sequences with varying lengths and similarities. (D) Multiple sequence alignment by Clustal Omega revealed that the first exon of SChLAP1 consists of three repeat regions, including Repeat 1 (R1), Repeat 2 (R2), and Repeat 3 (R3) (50). (E) Multiz alignment (top) and FASTA sequence alignment (bottom) for each exon. Black- and blue-colored regions represent sequence conservation. (F) Graphical representation of conserved nucleic acid sequences of SChLAP1. Multiple highly conserved motifs were identified throughout the second exon: ‘TGGCCATCAATA’ from Nucleotides 1-12, ‘TTCTGAAAATGGCAGT’ from Nucleotides 14-29, ‘TATAGAAGCCACTCTCACC’ from Nucleotides 53-71 and ‘ATGCACCTGGAAGCAACA’ from Nucleotides 74-91. The height of each stack represents the sequence conservation.

**Supplementary Figure S3.**
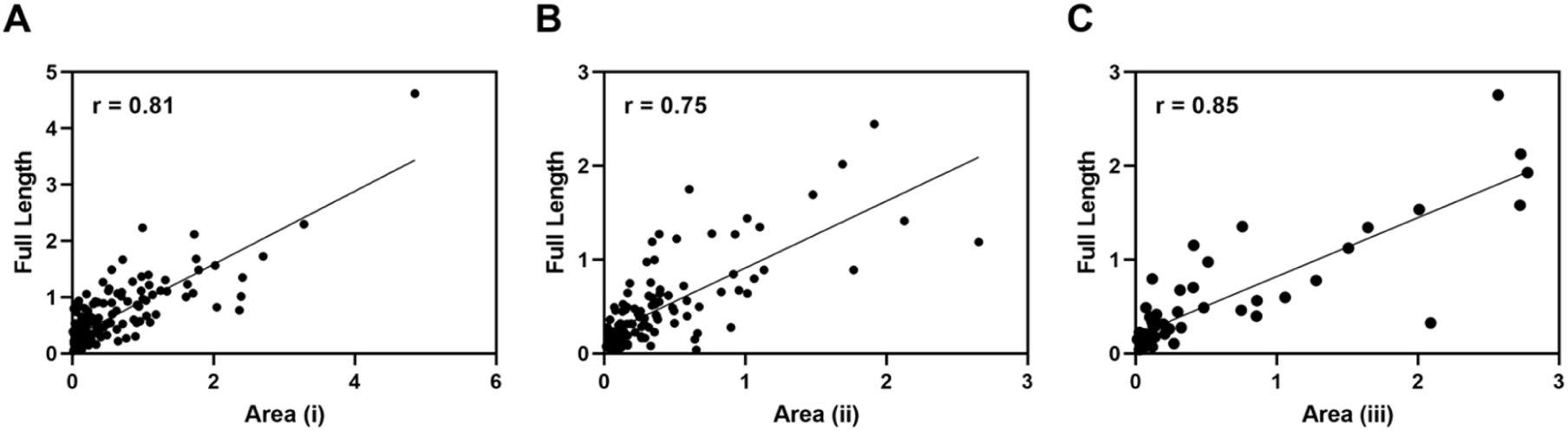
(A-C) Scatter plots comparing SHAPE reactivities of Area (i) through (iii) (see Figure 5) with the corresponding region in the full-length SChLAP1. The data represent the average of two biological replicates. Pearson correlation coefficient (*r*) values are indicated.

**Supplementary Figure S4.**
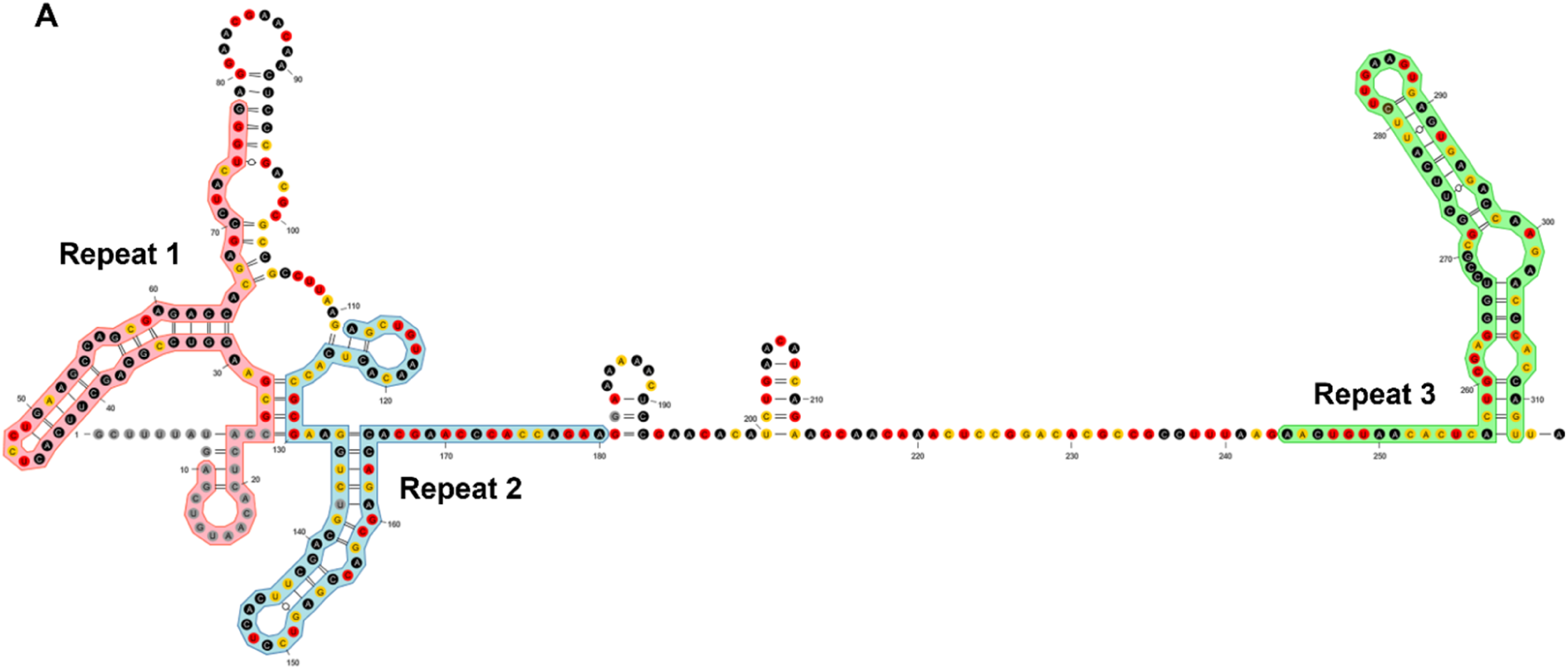
Sequential and structural analysis of the repeats. Each of the three repeats is highlighted using different color schemes.

**Supplementary Figure S5.**
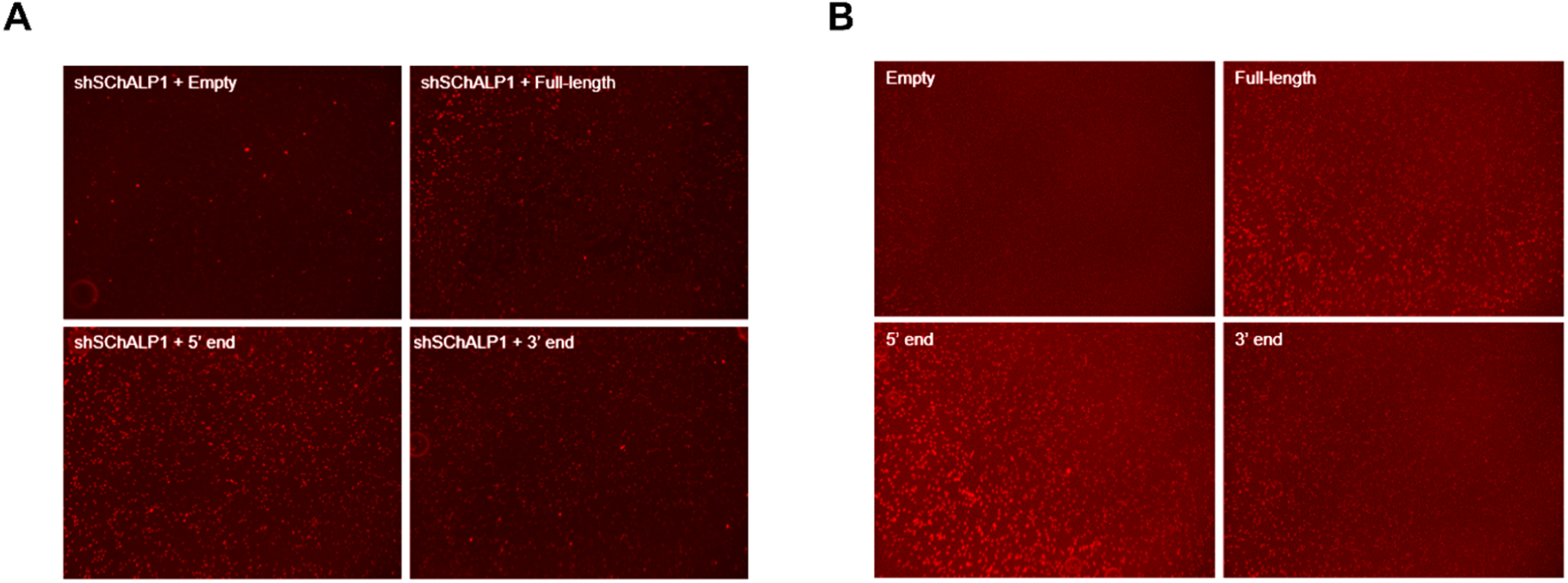
Representative images of transwell invasion assays in LNCaP (A) and 22Rv1 cells (B). Cell invasion was assessed by averaging data from three randomly selected fields.

**Supplementary Figure S6.**
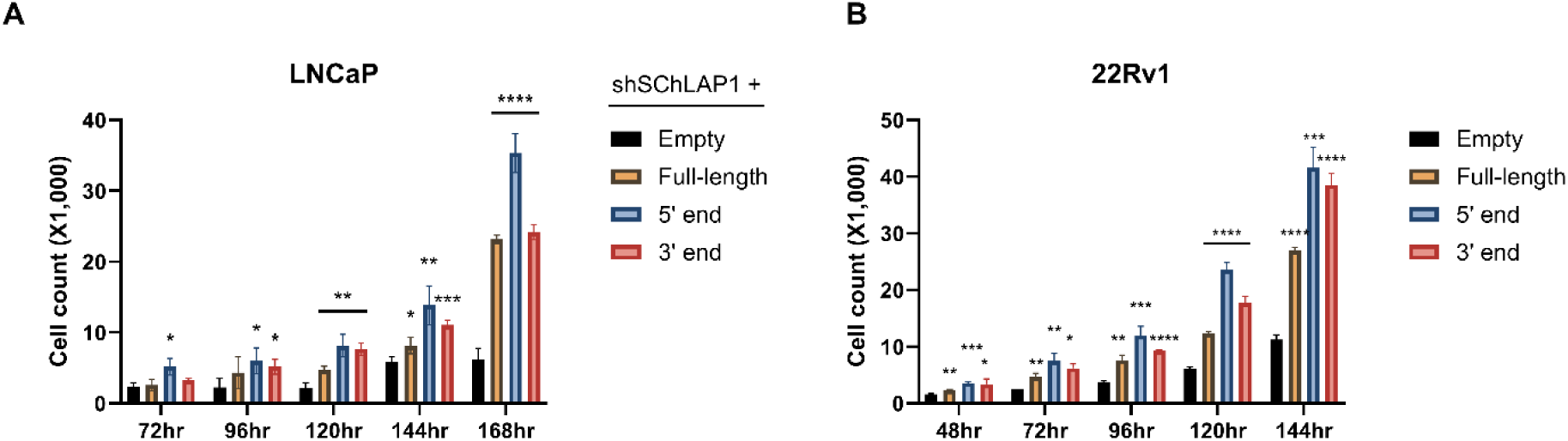
Cell proliferation was measured by counting cells upon overexpressing the 5’ or 3’ end of SChLAP1 in LNCaP (A) and 22Rv1 cells (B). These plots are bar graphs depicting the data shown in Figures 6H and 6I.

**Supplementary Table S1.**
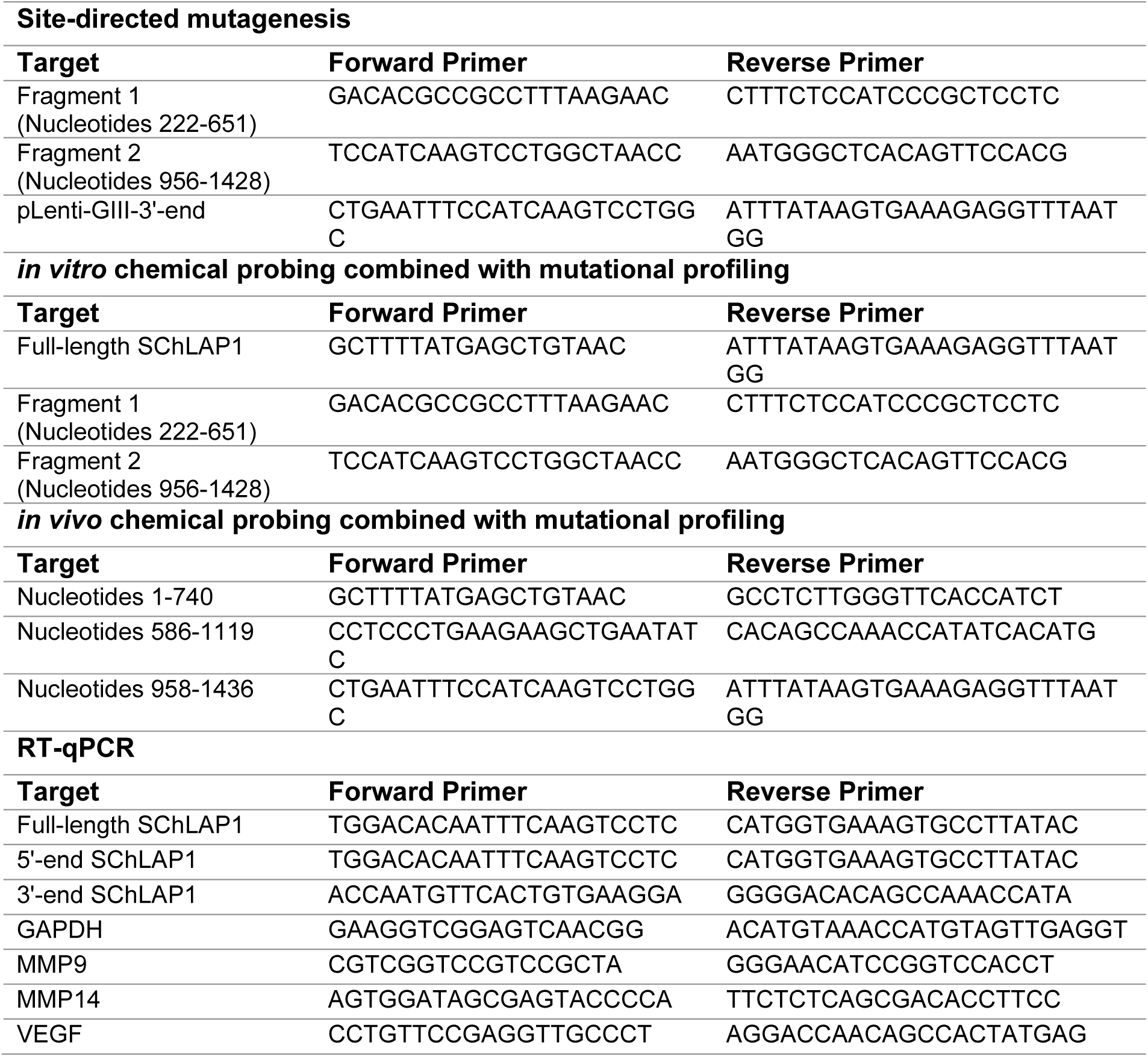
Primer sequences used in this study.

